# Prohlatype: A Probabilistic Framework for HLA Typing

**DOI:** 10.1101/244962

**Authors:** Leonid Rozenberg, Jeff Hammerbacher

## Abstract

HLA typing from sequencing data is considered as a classical probabilistic inference problem and Profile Hidden Markov Models (PHMM) are motivated for the likelihood calculation. Their generative property makes them a natural and highly discernible method; at the cost of considerable computation. We discuss ways to ameliorate this burden, and present an implementation https://github.com/hammerlab/prohlatype.

## Introduction

The major histocompatibility complex (MHC) is a set of genes found on the short arm of chromosome 6[19] that determine how antigen is presented to T-cells. In humans, this region has historically been referred to as the HLA (human leukocyte antigen) region. The variants of these genes, alleles, have various associations with disease as they modulate how our immune system behaves. HLA typing is the process of deducing these alleles from data.

Aside from disease association, accurate HLA-typing is valuable to medical procedures and future treatments. Matching compatible HLA types is important to avoid transplant rejection or specific complications such as graft-versus-host disease. Furthermore, precise HLA types are necessary for MHC binding prediction which has direct implications for immunotherapy.

### 1.1 Complexity

Accurate HLA typing is complicated by three main concerns; the genes of interest are homologous, they are highly polymorphic, and we have limited sequence knowledge of their variants.

Genes in this region are categorized into three classes: I, II, and III. Class I genes encode MHC molecules that are generally used to present peptides of non-self proteins to T-cells. Commonly, only three class I genes are considered: A, B and C. Although there are three other genes (E, F, G) and 14 pseudogenes (H, J^1^, K, L, N, P, S, T, U, V, W, X, Y and Z)[3] that complicate accurate typing.

Class II genes encode similar MHC molecules that, generally, present self peptides to different T-cells. These genes work in pairs to construct the final MHC protein and consequently the names of the genes have an A or B to determine the pairing: DMA, DMB, DOA, DOB, DPA, DPA, DPB, DQA, DQB, DRA, and DRB. Finally, Class III genes encode components of the complement system, and are sufficiently different to be excluded from analysis.

These genes exhibit strong sequence homology. An examination of the Levenshtein distances for exons of the class I genes (Table 1) shows that the distance across genes can be smaller than the distance within. For example, there are no differences out of 270 bases that comprise the second exon *B*:40:110* and *C*:16:85*. Similarly consider that there are only 12 changes between *A*30:02:05* and *H*02:04* (a pseudogene) in exon 3, whereas the farthest alleles are 22 and 8 bases different within their respective genes. This poses a problem for traditional alignment techniques for short reads in this region. As the database of reference sequences grows, the underlying indexing scheme in different aligners associates more and more alleles with each substring[12]. Aligners do not use the full, kilobase long, sequence as the target. Nor do common, short-read, sequence technologies span the full gene.

**Table 1:** HLA Class I homology, comparing Levenshtein distance within and across genes.

| Exon | Allele 1 | Allele 2 | Minimum distance | Farthest to Allele 1 | Maximum distance | Farthest to Allele 2 | Maximum distance |
| --- | --- | --- | --- | --- | --- | --- | --- |
| 1 | B*07:294 | J*01:01:01:01 | 6 | B*27:13 | 10 | J*02:01 | 2 |
| 1 | B*15:428 | K*01:01:01:01 | 6 | B*81:01 | 7 | K*01:02 | 1 |
| 1 | B*40:10:01:01 | L*01:01:01:01 | 4 | B*07:294 | 7 | L*01:01:02 | 1 |
| 2 | B*40:110 | C*16:85 | 0 | B*07:13 | 31 | C*04:01:02 | 38 |
| 2 | B*54:35 | G*01:01:01:01 | 27 | B*57:71 | 30 | G*01:01:12 | 2 |
| 2 | B*54:12 | H*02:04 | 13 | B*40:215 | 30 | H*01:02 | 7 |
| 2 | C*01:02:41 | E*01:01:01:01 | 31 | C*16:85 | 33 | E*01:01:02 | 1 |
| 2 | C*03:03:21 | G*01:01:01:01 | 27 | C*16:85 | 31 | G*01:01:12 | 2 |
| 2 | C*03:04:06 | L*01:01:01:01 | 31 | C*16:85 | 32 | L*01:01:01:02 | 0 |
| 2 | C*14:69 | Y*01:01 | 24 | C*16:85 | 33 | Y*02:01 | 2 |
| 3 | A*32:89 | B*27:71 | 9 | A*02:38 | 21 | B*07:78 | 22 |
| 3 | A*30:52 | C*12:96 | 13 | A*01:01:31 | 25 | C*17:19 | 17 |
| 3 | A*30:02:05 | H*02:04 | 12 | A*01:130 | 22 | H*03:01 | 8 |
| 3 | A*29:02:16 | Y*01:01 | 7 | A*02:38 | 18 | Y*03:01 | 0 |
| 3 | B*27:71 | A*32:89 | 9 | B*07:78 | 22 | A*02:38 | 21 |
| 3 | B*35:205 | C*15:117 | 2 | B*08:14 | 20 | C*04:77 | 15 |
| 3 | B*47:10 | Y*01:01 | 15 | B*08:84 | 22 | Y*03:01 | 0 |
| 3 | C*15:111 | B*40:120 | 2 | C*03:296 | 15 | B*08:14 | 19 |
| 3 | C*17:29 | H*02:04 | 17 | C*12:183 | 18 | H*03:01 | 8 |
| 4 | A*30:95 | Y*02:01 | 4 | A*02:411 | 14 | Y*01:01 | 13 |

For all class I genes as reported by IMGT, alleles where the full exon sequence is known were compared for minimum distance between genes. Subsequently, for both alleles of the minimum pair we looked for the maximum distance within their respective gene. The examples chosen maximal, atypical; they're purposefully chosen to demonstrate the parochial situations and to demonstrate the pervasive similarity amongst these genes. Exon distances are usually 73, 270, 276 and 276 base pairs, for exons 1, 2, 3 and 4. The full account of these distances is in Appendix A. Class II when compared against Class I does not have such similarities.

The homology problem is compounded by HLA’s extreme allelic heterogeneity. The canonical repository (IMGT[17]) reports several thousand alleles for multiple genes as of release 3.30 (Table 2).

**Table 2:** Number of alleles and their sources.

| Gene | A | B | C | DRB1 | DQB1 |
| --- | --- | --- | --- | --- | --- |
| Number of alleles | 3996 | 4858 | 3604 | 2120 | 1149 |
| GDNA derived | 648 | 740 | 772 | 83 | 165 |
| CDNA derived | 3348 | 4128 | 2832 | 2037 | 984 |
| Partial gDNA<sup>1</sup> | 1 | 1 | 0 | 0 | 0 |
| Partial cDNA | 3273 | 4021 | 2777 | 1960 | 942 |
| Relevant exon | 2310<sup>2</sup> | 2867<sup>2</sup> | 2178<sup>2</sup> | 1675<sup>3</sup> | 654<sup>3</sup> |

<sup>1</sup> Allele has missing data with respect to gene's reference sequence. <sup>2</sup> Exon 2 and 3. <sup>3</sup> Exon 2.

These genes are on the order of several kilo-bases long, implying a SNP frequency at least two orders of magnitude greater than the rest of the genome[5]. It is important to stress that our knowledge of this region is continuously changing as more alleles are ceaselessly reported to IMGT. This implies that *any* inference method based on the current dataset will report the wrong allele for a novel sample.

Finally, accurate typing is further complicated by our limited sequence knowledge of these alleles. Table 2 details the sparse reality of reference data. For over 50% of the Class I alleles we only have sequence information for exons 2 and 3 (out of 8)^2^. These may be the most relevant, as they determine the binding grove of the MHC molecule that presents peptides to the TCR[2], but unless our sequencing targets them specifically we have to take steps to impute the missing data or incorporate our lack of knowledge into the inference.

### 1.2 Existing Approaches

Existing tools, that target next generation sequence products, usually start with an alignment of reads to HLA reference sequences. Afterwards various procedures are followed to infer the alleles. Some tools focus on read and alignment quality and through multiple filters arrive at alleles (e.g. ATHLATES[13]). Other tools recognize that there is a natural constraint on the number of alleles that could explain the data; one allele per chromosome per gene. They formulate the inference as an optimization of which set of alleles account for the largest amount of the read data (e.g. OptiType[18], xHLA[20]). Yet other methods seek to creatively realign reads to a graph reference and deduce the alleles based on a likelihood score (e.g. HLA-PRG[8]). While these methods can be efficient and powerful, they have several limitations.

First, the scores that these models use for comparison or the constraints they try to optimize make errors into model errors; they encode an assumption about the distribution of read-data where the inference worked. If a user’s read data doesn’t match that distribution (e.g. different sequencing technology or number of reads), how does that tool’s model perform? This is a particularly pertinent for HLA inference for the reasons previously described. For specific examples consider that some tools make an arbitrary choice of selecting the second allele (of a diploid pair) based upon numeric limits. This means that zygosity is not modeled in the inference. Other tools will impose strict filters on which reads to use. Consequently, if there are reads which do not align well, perhaps from an unobserved allele, we would discard this evidence. Many tools focus on the “core” exons of genes (2 & 3 for Class I), but this makes a biased choice about the underlying distribution of read data that may be inappropriate.

Second, arbitrary score functions are arbitrary; they are not interpretable and do not have well understood semantics such as probability. Consequently, these methods make no allowance for incomplete inference, it may be the case that we do not have sequencing data at a specific position to differentiate between two alleles.

Lastly, there is limited or no description of the uncertainty in the inference. Some tools make a limited attempt to model the alignment (e.g. Seq2HLA [7]) as a normal distribution and compute statistics, but that is a choice based on modeling convenience and not necessarily a rigorous choice for a null-model.

## Results

### 2.1 Desired framework for inference

For the purpose of this framework our source of patient data are reads from next generation sequencing technologies. They provide a high throughput, but noisy channel of a patients genome. The data may originate from different sources (DNA, RNA) and may be targeted to specific parts of the genome, such as exome sequencing.

We abstract a read, *R*_*j*_, as an array of bases *R*_*j*_=*b*_*1*_ *b*_*2*_…*b*_*K*_, *b*_*i*_ *∈{A, C, G, T}* with an associated array of errors *ε*_*1*_ *ε*_*2*_ *ε…ε*_*K*_ that represent the probability that the reported base is *wrong*.

A simple way to perform HLA-typing is to compute the posterior distribution of HLA types;

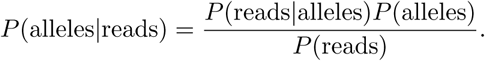

We’ll use our background knowledge in order to have precise, computable, equations for these values.

For a given gene (e.g. HLA-A) let *A*_*k*_ denote the set of alleles. At the outset, let us acknowledge that humans are diploid and our inference must provide a posterior over pairs of alleles, (*A*_*i*_ *, A*_*j*_), where *i* = *j* denotes a homozygous phenotype^3^.

The prior probability (*P*(alleles) = *P* (*A*_*m*_, *A*_*n*_)) should reflect a practitioners previous knowledge of allele distributions. We may lookup values from `allelefrequencies.net`, and use the observed human population frequencies or use knowledge of a patients ethnicity to provide a tighter set of values. But in practice we can be agnostic about this value, and assume a uniform probability, as we hope to have enough sequence information and consequently a discriminative likelihood (*P*(reads|alleles)) to overwhelm the prior and determine a posterior distribution with most probability mass in only one diploid.

The evidence (*P*(reads)), the probability of observing the reads that we have for our analysis serves as a normalizing constant for the product, so that our result is a probability.

When we consider our data, the reads, are presented in a specific order (*R*_1_, *R*_2_…*R*_*n*_), but for the joint read probability (*P*(reads) = *P* (*R*_1_, *R*_2_…*R*_*n*_)), the order of the reads, *R*_*i*_, should not matter, they are *exchangeable*[6]. Appealing to DeFinetti’s Theorem, they are identical, independently distributed, conditioned on the parameters that generated them; alleles, *A*_*k*_[15]. Consequently we can compute the likelihood as a product over individual reads,

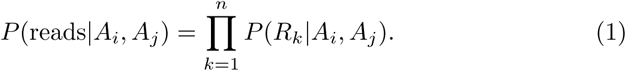

Our likelihood is conditioned on pairs of alleles, but a read almost certainly^4^ originates with only one of the two alleles. Therefore, for our likelihood, it would be principled to choose the higher probability

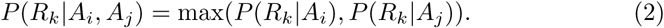

Lastly, to deal with the homology of this region we need to infer the types for all of the class I genes at the same time. The previous across and within gene distance survey (Table 1) should convince us that if *B*_*j*_ denotes the set of alleles for HLA-B and *C*_*k*_ the alleles of HLA-C, it is possible to construct (and encounter) a read where

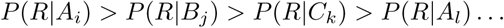

Consequently, we’ll choose the gene origin of a read based on the same *max* function

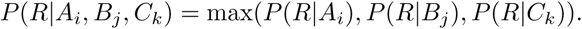

In practice this evaluation is extended to all class-I genes.

### 2.2 A powerful likelihood

We have reduced the structure of our inference to one remaining function; the conditional probability of observing a single read given an allele sequence. Part of the art (and power) of Bayesian modeling is choosing an appropriate likelihood function. Fortunately, a good candidate is already available in the bioinformatics community, Profile Hidden Markov Models (PHMM)[9].

HMMs represent sequential observations as emissions of hidden states. Probability distributions are used to determine the chance of an observation at a given hidden state, and transitions between the hidden states[16]. Traditionally, a “hidden” state models a condition that is difficult to determine precisely from the observation sequence. But they can have interpretable semantics as well, and an adroit choice of transition probabilities (ie. by assigning 0 to some) can serve to precisely structure the data generating process. Such is the case with our Profile HMM and reads from NGS.

Profile HMM were first considered in the context of multiple sequence alignments to help determine phylogeny. In this case, individual alignment positions had a (“hidden”) match state with an emission probability distributions to model conservation or alteration. In addition, insertion and deletion states between positions to handle gaps in the alignment.

We can measure the quality of a sequence’s alignment against a specified HMM model, by computing the probability that the model would output that sequence. This is usually accomplished with a *forward* pass. The “forward” comes from traversing the observations in the order that they appear. Computing this pass is a traditional dynamic programming problem, similar to the traditional string matching algorithms such as Needleman-Wunsch. But with an important caveat, as opposed to having different scoring options for matching, opening or extending gaps; we have a probability distribution over transitioning to a match, insert or delete state. When we aggregate emission from, and transitions between these states, we can assign a probability to the emission of a full sequence from an HMM. This is the likelihood function we seek.

We take inspiration from [11] for our formulation of the PHMM structure, which we will quickly summarize here but save a full description of our modifications for the section below where we discuss the computational burden. There is a *start* state *(S)*, that initializes where a read may match the reference. For each read position *l* (1 ≤ *l* ≤ *L)* and reference position *k* (1 ≤ *k* ≤ *K)* there are three states: *match* (*M*_*lk*_), *insert* (*I*_*lk*_) and *delete* (*D*_*lk*_). The *match* states emits (*e*_*M*_) the read’s base with the inverse probability of the error (*1 – ε*_*l*_) if it matches the reference and (*ε*_*l*_*/3)* it doesn’t.

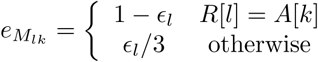

The *insert* states emits one of the four possible bases with equal probability, while *delete* states do not emit any symbols. There is an *end* state *(E)* for transitioning after the final read symbol. The *start (end)* state may emit a token symbol to represent the start (end) of a read; but the transition probabilities from (to) this state equally weigh the starting positions for possible paths that the read may take through the reference. The likelihood that we seek is the final probability of emission at the end state, equation 4. To obviate multiple levels of subscripts, we will use the symbol of the state to stand for the forward probability at that state.

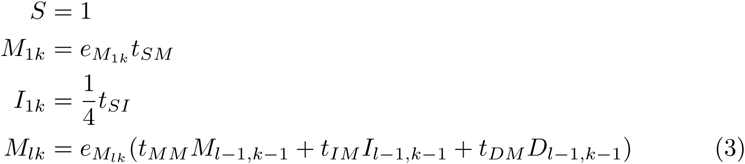

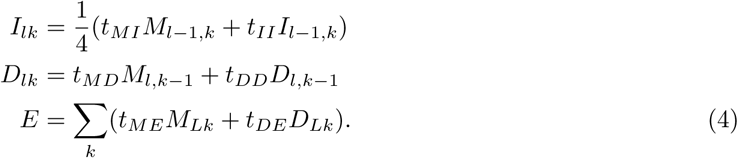

We’ll use transition probabilities (they are indexed by the origin and then destination states; e.g. *t*_*SM*_ = (1 − *α)*/*L)* similar to [11]

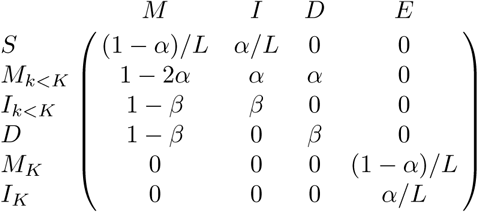

to further constrain the possible path through the HMM and to normalize the final likelihood of alignment. We use constants *α* = 0.001, which corresponds to probability of transitioning to the *insert* or *delete* state from the *match* state (this is similar to a gap opening penalty), and *β* = 0.1, the probability of staying in an *insert* or *delete* state (extending a gap)^5^.

We would like to note how this model closely resembles how one could generate a sequence from a reference. Suppose that some thing^6^ is at position *k* of a reference sequence having already made *i* characters of the sequence. It could proceed on to the next position and take the character from the reference, a match with a correct emission. It could proceed to the next position and make a mistake in the chosen character from the reference, a match with a incorrect emission. It could pause at position *k* and emit a random character, an insert. Or, lastly, it could skip that position entirely and move to *k* + 1, a deletion. We could think of other actions^7^, but until we observe those actions in our data, we can assign them a tiny probability, or, for the sake of computation, exclude them from our model. The point stands that if this process faithfully depicts how our data is generated, we should give more credence that it is correct than an arbitrary score.

### 2.3 Computational Cost

While performing the forward pass, for a read of length *L* and an allele of length^8^ *K*, we need to compute three values (*M*_*lk*_*, I*_*lk*_*, D*_*lk*_) for each *l* and *k* position. This can be accomplished recursively in *O(KL).*

Once we factor in that we have *M* reads per sample and *N* alleles, we’re faced with a potential *O(KLMN)* total cost to type one patient. There is no denying that any realistic assignment of these variables (*L* = 100, *K* = 4000, *M* = 1000 and *N* = 3000) leads to an almost infeasible problem for ordinary computers. Perhaps this is the reason why PHMMs despite their pervasive use in bioinformatics have not been used to address this problem. Are there techniques that would allow us to pare down the steps necessary to perform this calculation? Yes, our main contribution is that we can replace *N* with a small constant.

#### 2.3.1 Embedding

To replace *N* consider the multiple sequence alignment of the alleles. Conveniently, IMGT provides such an alignment that we use as reference. At each position of this alignment, each allele can be in one of 6 states; 4 nucleobases, gap and missing. In the alignment, the first listed allele is the reference allele of the gene (e.g. “A*01:01:01:01”). The gap character in the reference allele indicates that some other alleles (listed below) have an insertion of bases. Whereas a gap character for the non-reference alleles indicates a deletion of bases from that allele, with respect to the reference. We’ll discuss later how we handle missing positions, but for current purposes, assume that we will map these positions to a nucleobases from another allele (such as the reference). This leaves us with only 5 states and consequently, for a specified base from the read, there can be at most 5 possible emissions of the PHMM. Therefore, for one read, we can calculate the probability of emission for all of the alleles with one pass if during every calculation we maintain a mapping of alleles to state probabilities. As long as the number of state probabilities that we track is smaller than the number of alleles, we can be faster than *O(LKN)*, per read.

There are a couple of modifications to naively computing the PHMM in order to be able to calculate the emission for all of the alleles with one pass. The most important is a representation of the states for all of the alleles simultaneously, we will discuss an efficient structure for this momentarily, but for the moment assume that we have some ordering of the alleles (*A*_1_, *A*_2_…*A*_*n*_…*A*_*N*_) and that we can extend each state to take a third index for the allele: (e.g. *M*_*ikn*_ is the probability of the *n*th allele being in the *match* state after seeing *i*th base of the read and the *k*th alignment position. But notice that in the alignment, there might *not* be such a base; the *n*th allele might be in a gap. Consequently, we need to rewrite our forward equations such that they incorporate the gaps. Let us define an indicator variable

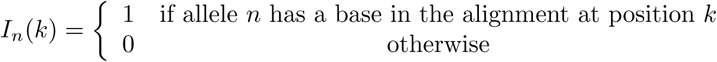

and *p*_*n*_*(k)* be the first position < *k* in the *alignment* where allele *n* has a base (for the majority of case this is just *k* − 1). This leads to the following forward equations:

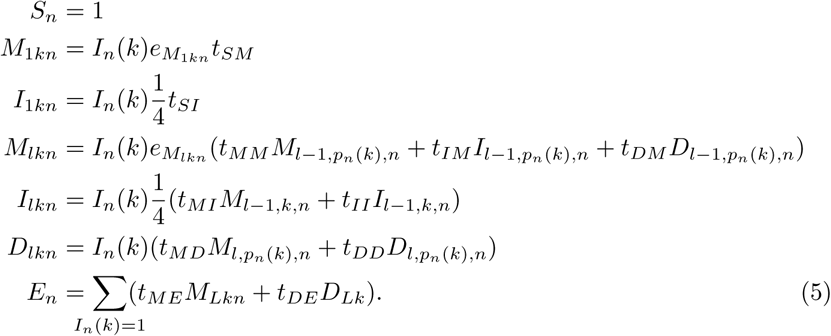

Even though, for a given state and allele, we can define a zero forward probability via the indicator *I*_*n*_*(k)*, that is not how we want the read to align against that allele. The global alignment is a convenient construct, not how we want to model the data generation of the reads, and hence we use *p*_*n*_*(k)* to index around the gaps in the alignment An equivalent way of formulating these recursions without needing to explicitly keep track of *p*_*n*_*(k)* is to pass the delete state *D*_*l* − 1,*k* − 1_ for alleles that do not have an emission at *k*.

#### 2.3.2 Partition map

If we keep track of all *N* states for each of the *M*_*lk*_, *I*_*lk*_, *D*_*lk*_ then we are back into the troublesome *O(KLMN)* total running time. But the previous equations make it easier to observe that if allele *n* and *n* + 1 have the same base at *k* (ie. *A*_*n*_ *[k] = A*_*n* − *1*_ *[k]*), then they’ll have the same emission *e*_*M* *lkn*_ and consequently the same first match state (*M*_1 *kn*_). And their forward states will continue to be equivalent up to the point where they may^9^ disagree.

This suggests the second necessary modification, rather than using a linear data structure such as an array to store the forward values, per allele, we will use a compressing structure similar to a run-length encoded list. We will save the full description of this data structure for appendix B but it is important to note two properties. First, the resulting data structure has size roughly proportional to the number of unique values stored therein, *u << N*. We say roughly because the keys used to index these values are compressed form of representing the alleles, where similar alleles (based on values) are stored next to each other. Combined, this accounts for the good performance.

#### 2.3.3 Details

The order of the alleles matters a great deal. Randomly ordering alleles will slow down the merging, whereas the order according to the HLA nomenclature[4] has worked pretty well.

Our final improvement is to bind the precision of floats in the calculation. A sensible implementation of HMMs will use 1 of 2 techniques to remain accurate. The first would normalize the probabilities at fixed *i* to prevent the values from overflowing. The second approach would use *log*-probabilities throughout the calculation and only return to probabilities (and normalize), at the end. This is doubly convenient as error probabilities in the FASTQ format are already reported in log_10_format. Since the smallest quality score is 6 × 10^−6^ we constrain the resolution of the oats to 1 × 10^−6^. This prevents numerical inaccuracies from creating values that are pointlessly different, which helps to constraint the previously mentioned *k*.

### 2.4 Missing reference data

PHMMs provide an work-around solution to incomplete allele sequence data.

For an allele *A*_*m*_ that is missing a region *R*, we can borrow it from an allele with that region *A*_*p*_. Notice that only equation 3 depends on an alleles base position via the emission (*e*_*Mlk*_). Therefore when we borrow bases (or gaps) from *A*_*p*_ for all *k∈R*, *M*_*lk*_ (*A*_*m*_ *) = M*_*lk*_ (*A*_*p*_). Consequently, *P*(*R|A*_*m*_) will have the right relationship with respect to *P(R|A*_*p*_). To the extent that *P(R|A*_*m*_)≠*P*(*R|A*_*p*_) it will be determined by the sequence information we have, outside the missing region *R*.

Unfortunately, this relationship does not extend cleanly to multiple alleles. Consider without loss of generality a third allele *A*_*q*_ that is not missing data from region *R*, but we will still borrow from *A*_*p*_. We’re only interested in the behavior in *R*. As described *P(R|A*_*m*_ *) = P(R|A*_*p*_), because *A*_*m*_ is borrowing *A*_*p*_’s sequence information. *A*_*p*_ and *A*_*q*_ have data and by construction the more likely allele will have the higher probability and so the likelihood of *A*_*m*_ will be in line with *A*_*p*_. But what if, unbeknownst to us, *A*_*m*_’s real sequence is more similar to *A*_*q*_ then *A*_*p*_ in *R*. Then reads aligned there will be incorrectly aggregate in the results (equation 1).

In practice this limitation is not that troublesome. There is more variation in regions where we’re not missing data (Table 3), and consequently the likelihoods calculated in that region are more discerning. If we have fairly even distribution of reads, these tend to out weight potentially bad choices for the imputation allele. We’ll address this issue further in the implementation.

**Table 3:** Variations per 100^1^ bases in Exons.

| Number of alleles<sup>2</sup> | A | B | C |
| --- | --- | --- | --- |
|  | 772 | 847 | 895 |
| Exon 1 | 30.14 | 32.88 | 28.77 |
| Exon 2 | 67.04 | 51.48 | 68.15 |
| Exon 3 | 71.01 | 55.43 | 81.52 |
| Exon 4 | 25.36 | 20.29 | 27.90 |
| Exon 5 | 18.80 | 23.08 | 27.50 |
| Exon 6 | 21.21 | 9.09 | 18.18 |
| Exon 7 | 6.25 | 15.91 | 12.50 |

<sup>1</sup> Counting only SNV, not gaps, from the reference sequence.
<sup>2</sup> Selecting alleles where we have full sequence knowledge for exons 2 to 6.

### 2.5 Splitting

Lastly, as we iterate down the read and fill in the matrix that keeps track of the previous emission values, the variation in values between alleles increases monotonically; each alignment position is a potential point of disagreement. Consequently, the time to process a row corresponding to one read position also increases monotonically, and the last positions are the most costly to process. Fortunately, we can split the read. For a read *R* let 
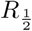
 be the first half and 
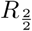
 the second half then

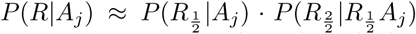
 and we have a natural divide-and-conquer strategy to speed up the calculation. Perform the forward pass on the first half, then on the second half and finally multiply the results.

The approximation comes from the in-exact modeling of the sequence generation. As opposed to constraining the transitions of the before mid-point base directly to either a *match*, *insert* or *delete* state; we’re transitioning to an *end* and then another *start* state first, which are averaged over all reference positions. The averaging introduces a non-linear transition as we now consider paths that are potentially backward!

## 3 Implementation

We’ve implemented the logic discussed in OCaml[10]. The source code is available at https://github.com/hammerlab/prohlatype. The project is a collection of tools that allows the user to calculate the HLA likelihoods of their read data, collectively referred to as “prohlatype”.

### 3.1 Protocol

The workflow to arrive at these HLA types consists of:

1. Download the IMGT-HLA database. The canonical repository is now easily available at https://github.com/ANHIG/IMGTHLA.
2. Create an imputed HLA reference sequences via the tool **align2fasta** found in the prohlatype project. This tool uses various imputation logics to extend allele sequences provided by IMGT to cover their locus.
3. Use the previously generated sequence database to filter one’s data via an aligner such as razers3 or bwa. Reads that align poorly to the HLA region will have a low likelihood for any gene or allele, they will act as a background noise in the distribution. Unfortunately, per-read inference, in prohlatype is still an expensive operation, and consequently it is more effective to filter your data to a smaller set of data. One can use lax filter parameters to form a more inclusive set of read data.
4. Export the reads from the filtered bam file to a FASTQ (via samtools) file.
5. Finally, one can calculate the likelihood with **multi_par**. This tool takes as input the resulting read les from (step 4), and a specification of HLA gene alignment from the previously downloaded IMGT-HLA database (step 1).

### 3.2 Output

**multi_par** supports two main output modes, tab delimited and JSON. Regardless of format the output is divided into two main sections. At the beginning, for each gene, there is a “Zygosity,” and “Likelihood,” section. Afterwards a “Per Read” section gives individual per-read assignments.

#### 3.2.1 Zygosity

This section gives the diploid *log*-likelihoods and normalized posterior probability^10^, lower numbered alleles are chosen as “Allele 1”.

| Allele 1 | Allele 2 | $\log_{10} P$ | Prob |
| --- | --- | --- | --- |
| A*23:17 | A*30:114 | -7096.62 | 0.12396 |
| A*23:17 | A*30:01:01 | -7096.62 | 0.12396 |
| A*23:01:19 | A*30:114 | -7096.62 | 0.12396 |
| A*23:01:19 | A*30:01:01 | -7096.62 | 0.12396 |
| A*23:01:01:03 | A*30:114 | -7096.62 | 0.12396 |
| A*23:01:01:03 | A*30:01:01 | -7096.62 | 0.12396 |
| A*23:01:01:01 | A*30:114 | -7096.62 | 0.12396 |
| A*23:01:01:01 | A*30:01:01 | -7096.62 | 0.12396 |
| A*23:01:01:02 | A*30:114 | -7098.10 | 0.00412 |
| A*23:01:01:02 | A*30:01:01 | -7098.10 | 0.00412 |
| A*23:01:01:02 | A*30:106 | -6890.22 | 0.00000 |
| A*23:17 | A*30:106 | -6893.32 | 0.00000 |
| ... | ... | ... | ... |

We interpret this to indicate that for one of the chromosomes there was almost equal evidence for 4 alleles (A*23:17, A*23:01:19, A*23:01:01:03, A*23:01:01:01) and less evidence for another allele (A*23:01:01:01:02). While the evidence for the other chromosome was evenly divided between 2 alleles: A*30:114 and A*30:01:01. Note that because of the variation in the second set of digits (17 vs 01 and 114 vs 01), this implies that for both chromosomes we’re uncertain of the final protein.

In this particular instance, subsequent analysis, by looking in the *per-reads* section, can explain this distribution. If we inspect the reads aligned to the first chromosome, we will find that there are only 5 reads aligned to the intronic region where there is a SNP accounting for A*23:01:01:02^11^ and no reads at the position inside of the 4th exon accounting for A*23:17. Similarly, for the second chromosome, there no reads covering the SNP in the 4th exon accounting for A*30:114.

Detailed analysis of why the distribution takes its shape requires further tooling, but knowing that all of the probability mass is not concentrated in one pair, or, more importantly, is distributed across alleles with different exonic sequences should inform clinical decisions.

#### 3.2.2 Likelihood

This section is a table where each row corresponds to one of the alleles in the locus that are considered for typing. For each allele we report the aggregate *log*_10_ likelihood of the read data^12^ and a description of the imputation steps taken to construct the full sequence for that allele. By themselves, the *log*-likelihoods can only serve as a comparison for the diploid likelihood in the *zygosity* section.

#### 3.2.3 Per Read

Here we detail how each read was aligned against all the loci under consideration. In particular, which orientation (regular or reverse complement) was chosen as most likely and if any filters were used to expedite computation. Furthermore, it describes the most likely emission position for the loci against which the read was matched. This information can later be aggregated to explain some non-descript zygosity descriptions.

## 4 Results and Discussions

Validation remains a considerable challenge for this work. While our observations of the world always form some empirical distributions, how we derive that distribution from different data sources requires care. If an oracle were to specify class I HLA types, we would expect 6 allele names, 2 for each of HLA-A,B,C. We would interpret this as a probability distribution with 1 assigned to this 6-tuple^13^ and 0 to all other allele combinations^14^.

The work in this project represents one framework for constructing such a distribution. But for comparison, how would we evaluate against other HLA-typers? Most tools do not provide measures of uncertainty with their assignments, nor do the reported p-values constitute an easily interpretable null-model that one could translate. We may treat them as oracles, but only to interpret their output as assigning all of the empirical probability mass to their chosen 6 tuple, not as being a source of truth. Finally, it needs to be noted that metrics on distributions (such as KL-divergence) are not easily interpreted as they require context.

Consequently, our analysis of concordance, between other HLA tools and prohlatype, aims for simple interpretation. First, we divide the analysis by loci. For each loci we will further divide the per sample interpretation into three categories based upon how many of the two alleles have considerable^15^ probability mass according to prohlatype: “both match”, “one match” and “both different.”

### 4.1 Concordance with 60 patient UPenn dataset

For a sample 60 patients who were previously exome sequenced for study on the genetic relationship of atopic dermatitis[14], we receiving HLA types determined via Omixon’s “Target” software. We typed these samples with OptiType and there was high concordance with the Omixon results. We ran prohlatype following the protocol specified in subsection 3.1 and compared the results. The reported types were to 4-digit resolution (e.g. A*01:01), so we aggregated all of the probability of alleles where those digits matched.

As we can see from table 4, usually more than two-thirds of the time Omixon reported a diploid pair to which prohlatype assigned probability. Given that, we can ask what probability mass is unexplained, and unfortunately observe that between 20% to 55% is missing. For the other samples where either one or both of the alleles are not present, the unexplained *P* is 100%. It is also important to note that all cases of claimed homozygosity by Omixon or OptiType were not substantiated by prohlatype.

**Table 4:**
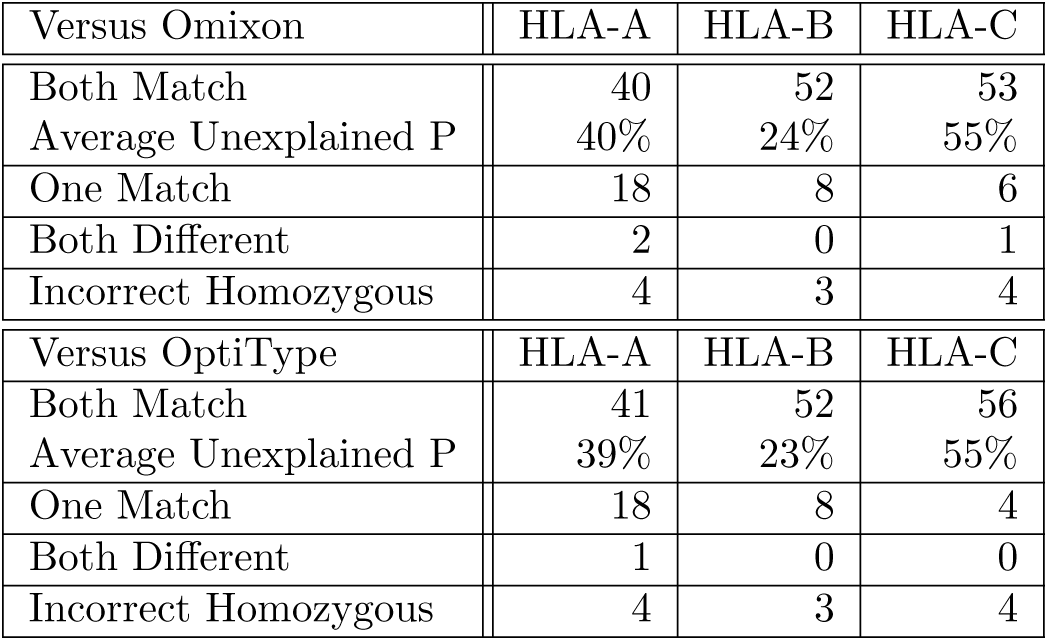
Concordance with Omixon and OptiType validated data, *n* = 60.

### 4.2 Comparison against a commercial lab

For an independent personalized genomics vaccine trial, we obtained sequencing on 5 patients and HLA types from a commercial lab specializing in this inference. We used prohlatype and compared against the well regarded labs results. The commercial lab performed independent sequencing targeting various exons of the HLA genes and performed inference using their custom software. They reported HLA’G’ group annotations[1], which groups alleles that have the same coding region. Our sequencing was 125bp long, untargeted. Out of 30 alleles, prohlatype reported the same alleles in 28 cases. Prohlatype’s output has higher resolution since it specifies the most likely allele within the G-group, but there is no probability mass associated with a potentially different protein product. Because of the commercial lab’s high regard we investigated the differences, they are informative and merit a detailed description.

#### 4.2.1 Patient 5: C*05:01:01 vs C*05:111

The commercial lab reported C*05:01:01G, while prohlatype reported C*05:111^16^. In trying to resolve this ambiguity we noticed that these two alleles (C*05:01:01:01 as the representative sequence for this group), differ only a single point; 25 base pairs into the third exon (referred to as position *p)*, C*05:01:01:01 has an A while C*05:111 has a C. Ordinarily, small differences like this are what give prohlatype its resolution, but in this case we wanted to investigate further given the commercial labs report.

It turns out that Patient 5’s other HLA-C allele is C*08:02:01:01 (both reports agree), which has a particularly interesting property: at *p* this allele also has an A and 125 bases on either side of *p* (referred to as region *r)* it has an identical sequence to C*05:01:01:01.

When we further inspected our read data, out of 60 reads that align *r* 1 read actually had a C at *p* in favor of C*05:111^17^, while the other 59 reads all had an A, in favor of either C*08:02:01:01 or C*05:01:01:01.

We know that C*08:02:01:01 is one of the patients correct alleles because we have matching read data in other parts of the gene. But in this one region, this second allele, acts as a mask on the inference. Consider three scenarios of what might be occurring at *r*:

1. 59 reads with an A at *p* come from C*05:01:01:01 and 1 read with a C at *p* from C*05:111.
2. 59 reads with an A at *p* come from C*08:02:01:01 and 1 read with a C at *p* from C*05:111.
3. There is one read with an error, that mimics an existing allele, C*05:111. Some proportion of these reads come from C*08:02:01:01 and the remaining from C*05:01:01:01. Furthermore, we have no way of knowing from which strand these reads originate.

We have read data from outside *r* that makes us favor C*08:02:01:01 as one of the two alleles and consequently we can disregard scenario 1. Prohlatype considers all possible assignments and as a consequence of equation 2 scenario 2 is more likely. But does that match our intuition?

Ultimately, we have to answer in the negative. We believe that scenario 3 is more likely than scenario 2, because we have a prior belief that our read distributions will be balanced. Specifically, we think that the strand bias in our data won’t be too great, or at least it will be consistent across our gene. We have other information that we want to incorporate into our inference.

#### 4.2.2 Patient 5: A*26:01:01:01 vs A*26:08

The commercial lab reported A*26:01:01G while prohaltype reported A*26:08^18^. We describe this difference second because after exhaustive investigation, there does not seem to a good explanation of incorrect inference, or defects within the framework.

What is interesting about this example (why we chose to write about it) is that A*26:01:01:01 happened to be the second most likely allele for this strand according to prohlatype. Furthermore, the evidence for these two alleles seems to be very close. In our data, 120 reads provided evidence that favor A*26:08, while 111 reads provided evidence that favor A*26:01:01:01. Consequently, the inference may not be as sharp in log-probabilities as we desire, but when normalized the probability is decisively^19^ in favor of A*26:08.

Again, were it not for having independent validation of the HLA types from a highly reputable commercial lab we would not provide detailed investigation of per allele and per read assignments. We can only surmise that the vagaries of next-generation sequencing lead to these to disparate results.

### 4.3 Do we need to type all of class I?

To highlight the need to type all of the class I genes at the same time, consider that for the previously described UPenn data set, 15% of all reads, that were previously aligned for only HLA-A,B,C with razers3 were actually for HLA-H.

**Table 5:**
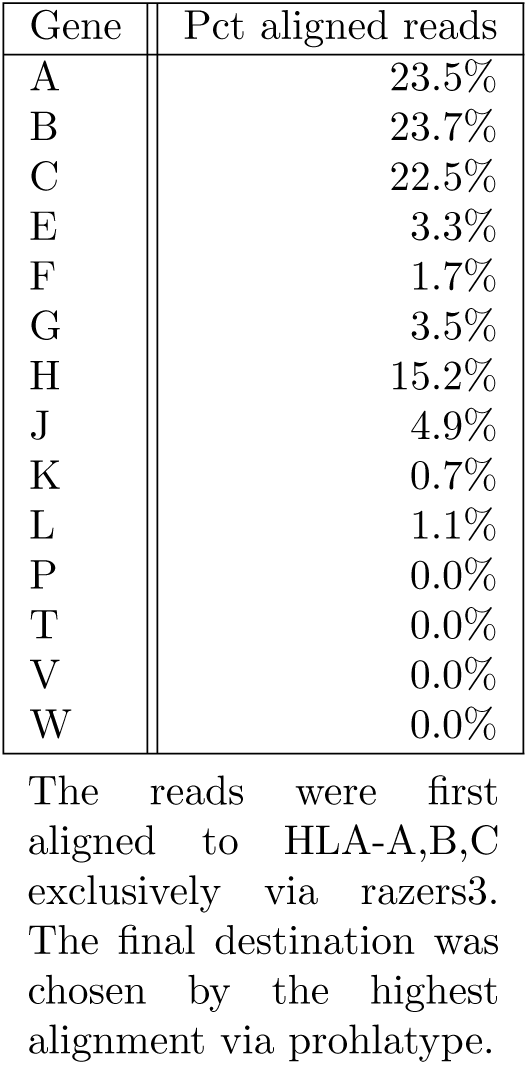
Percentages of reads aligned to genes.

| Gene | Pct aligned reads |
| --- | --- |
| A | 23.5% |
| B | 23.7% |
| C | 22.5% |
| E | 3.3% |
| F | 1.7% |
| G | 3.5% |
| H | 15.2% |
| J | 4.9% |
| K | 0.7% |
| L | 1.1% |
| P | 0.0% |
| T | 0.0% |
| V | 0.0% |
| W | 0.0% |

The reads were first aligned to HLA-A,B,C exclusively via razers3. The final destination was chosen by the highest alignment via prohlatype.

**Table 6:** Parametric PHMM per read timing comparison.

| Gene | Parametric<sup>1</sup> | Projected<sup>2</sup> | Pct |
| --- | --- | --- | --- |
| HLA-A | 9.41 | 168.94 | 5.57 % |
| HLA-A | 8.08 | 168.00 | 4.81 % |
| HLA-A | 9.22 | 169.89 | 5.43 % |
| HLA-A | 7.69 | 169.97 | 4.52 % |
| HLA-A | 9.13 | 169.94 | 5.37 % |
| Average |  |  | 4.83 % |
| HLA-B | 9.51 | 218.12 | 4.36 % |
| HLA-B | 9.60 | 217.65 | 4.41 % |
| HLA-B | 9.14 | 217.19 | 4.21 % |
| HLA-B | 11.54 | 218.21 | 5.29 % |
| HLA-B | 9.52 | 217.28 | 4.38 % |
| Average |  |  | 4.50 % |
| HLA-C | 11.39 | 236.39 | 4.82 % |
| HLA-C | 7.16 | 236.10 | 3.03 % |
| HLA-C | 11.33 | 237.46 | 4.77 % |
| HLA-C | 10.72 | 237.70 | 4.51 % |
| HLA-C | 9.61 | 237.28 | 4.05 % |
| Average |  |  | 4.48 % |

Displaying 5 of 30 reads (per gene) used for analysis, average is over all 30. <sup>1</sup>Time, in seconds, to perform parametric PHMM forward pass. <sup>2</sup>Projected time, in seconds to perform PHMM forward pass for all alleles.

### 4.4 Performance

To document the improvements in performance we compare the time of one parametric PHMM pass versus the projected time to calculate the forward passes individually, one allele at a time. For HLA-A, B, and C we randomly chose 50 alleles and 30 reads, that were previously aligned to those genes. For each allele we timed (average of 10 times) how long it would take to perform a regular PHMM forward pass for each read. We compute the average time across the alleles, per read, for comparison. For each read we timed (average of 10 times) how long it would take to perform one parametric PHMM’s forward pass.

There are several factors that impact performance of parametric PHMMs and complicate an algebraic run-time analysis of the parametric maps.

1. The number of alleles.
2. The order of the alleles. We want similar alleles to be next to each other for better compression.
3. The position of the read. In area’s of lower allele variance the partition maps are smaller.

#### 4.4.1 Splitting

For the UPenn dataset splitting into 2 halfs reduces the average user CPU time by 30%, while splitting it into quarters reduces the time by 37%. It seems like there are diminishing returns to splitting further.

### 4.5 Choice of imputation allele

We investigated three strategies for choosing which allele to borrow from when we have limited sequence information.

**Reference:** Always choose the reference allele, which has sequence knowledge spanning the full alignment.

**Trie:** Choose the allele with the closest lexicographic name with available sequence knowledge. The allele name form a four level tree based upon their nomenclature[4], where each subsequent field in the names forms a node of the tree. For an allele missing data we would walk up the tree, starting with the last named field looking for an allele which has the missing region.

**Segments:** Group alleles by the segments, regions of their gene, where we have sequence knowledge. Then measure allele similarity with Levenshtein distance over those segments. Choose the imputation allele that is closest over these distances.

We’ve tested all three methods for the UPenn dataset. In practice they give almost identical results; they ascribe almost identical probability mass to the same alleles. The discrepancies occur when there is a paucity of data such that the final distribution is broad. In this case, numerical instability assigns insignificantly different final probability values. The trie method is slightly (10%) faster and consequently is the default method.

## 5 Conclusions

We have presented a framework for Bayesian inference of HLA-types and discussed an implementation that makes the computation feasible.

### 5.1 Class II

This project started as an investigation of HLA-typing for class II genes based on exome sequencing. But finding reliable models for class I motivated and presented new challenges sufficient for this body of work. Class II presents a couple of novel additional challenges. First the total alignment sequence is more than 10 times longer, this will make the total required computation time infeasible for all but the most patient of researchers. It merits new techniques to improve performance

Second, there are bigger structural gaps between different alleles that might complicate inference. Although, they might provide clearer resolution as well. This uncertainty introduces issues of how to correctly structure the computation. Akin to how the use of alternative class I genes (E, F…) improved the accuracy of typing, for class II there is a similarly large group of non-common genes (DRB3,DRB4, and DRB5) and pseudogene (DRB2, DRB6, DRB7, DRB8, DRB9), which provide variations for DRB1. This presents a challenge to figure out the right way to measure alignment against all of these loci.

### 5.2 Future work

There are three main thrusts of future work for this project. The first task is to develop an appropriate “regularization” framework to address issues where the dominance of equation 2 is mitigated. Equation 2 is a good choice for discerning an individual read’s likelihood given a diploid pair, but it sometimes leads to sub-optimal inferences in aggregate. We need to add our intuitions about strand balance into the framework to act as the regularizer.

The next step is to address novel allele discovery. All inference up to this point chooses the most likely alleles from those in the database. But, as we have acknowledged before, the set of alleles grows with time and as new and diverse human populations are sequenced it is not unlikely that we will attempt to infer a previously unobserved HLA-type.

We believe that PHMMs actually give an elegant algorithm for this discovery. Start with the alleles inferred using the methodology. For each read, compute the Viterbi pass, and inspect it to see if it has novel mutations, with respect to this most-likely. Add these mutations to a set of pseudo-alleles. Repeat this for all reads, and again using Bayes rule see if one of these pseudo-alleles has a higher posterior probability. We can use something like the Turing-Good approximation as a starting prior on these pseudo-alleles.

The remaining area of work, as previously mentioned, is to extend this to class-II genes.

## Acknowledgments

This research was supported by the Parker Institute for Cancer Immunotherapy. The authors are indebted to helpful discussions with Isaac Hodes, Sebastian Mondet and Arman Aksoy, and the other members of Hammer Lab.

## A Extended Homology Search

| Allele 1 | Allele 2 | Minimum distance | Farthest to Allele 1 | Maximum distance | Farthest to Allele 2 | Maximum distance |
| --- | --- | --- | --- | --- | --- | --- |
| A*03:01:58 | B*08:01:20 | 6 | A*02:01:132 | 4 | B*27:13 | 10 |
| A*03:01:58 | C*02:02:01 | 7 | A*02:01:132 | 4 | C*17:03:01:01 | 5 |
| A*01:01:01:01 | E*01:01:01:01 | 17 | A*02:01:132 | 3 | E*01:01:01:02 | 0 |
| A*01:01:01:01 | F*01:01:01:01 | 15 | A*02:01:132 | 3 | F*01:01:02:01 | 1 |
| A*03:01:58 | G*01:01:01:01 | 5 | A*02:01:132 | 4 | G*01:01:02:01 | 2 |
| A*03:01:58 | H*01:01:01:01 | 5 | A*02:01:132 | 4 | H*02:02 | 6 |
| A*03:01:58 | J*01:01:01:01 | 5 | A*02:01:132 | 4 | J*02:01 | 2 |
| A*03:01:58 | K*01:01:01:01 | 5 | A*02:01:132 | 4 | K*01:02 | 1 |
| A*03:01:58 | L*01:01:01:01 | 6 | A*02:01:132 | 4 | L*01:01:02 | 1 |
| A*01:237 | Y*01:01 | 1 | A*02:01:132 | 4 | Y*02:01 | 0 |
| B*08:01:20 | A*03:01:58 | 6 | B*27:13 | 10 | A*02:01:132 | 4 |
| B*15:428 | C*02:02:01 | 2 | B*81:01 | 7 | C*17:03:01:01 | 5 |
| B*40:01:01 | E*01:01:01:01 | 19 | B*27:13 | 8 | E*01:01:01:02 | 0 |
| B*15:428 | F*01:01:01:01 | 15 | B*81:01 | 7 | F*01:01:02:01 | 1 |
| B*07:02:01:01 | G*01:01:01:01 | 6 | B*27:13 | 9 | G*01:01:02:01 | 2 |
| B*07:02:01:01 | H*01:01:01:01 | 8 | B*27:13 | 9 | H*02:02 | 6 |
| B*07:294 | J*01:01:01:01 | 6 | B*27:13 | 10 | J*02:01 | 2 |
| B*15:428 | K*01:01:01:01 | 6 | B*81:01 | 7 | K*01:02 | 1 |
| B*40:10:01:01 | L*01:01:01:01 | 4 | B*07:294 | 7 | L*01:01:02 | 1 |
| B*08:01:20 | Y*01:01 | 7 | B*27:13 | 10 | Y*02:01 | 0 |
| C*02:02:01 | A*03:01:58 | 7 | C*17:03:01:01 | 5 | A*02:01:132 | 4 |
| C*02:02:01 | B*15:428 | 2 | C*17:03:01:01 | 5 | B*81:01 | 7 |
| C*03:03:36 | E*01:01:01:01 | 20 | C*07:02:80 | 6 | E*01:01:01:02 | 0 |
| C*02:02:01 | F*01:01:01:01 | 16 | C*17:03:01:01 | 5 | F*01:01:02:01 | 1 |
| C*03:02:01 | G*01:01:01:01 | 6 | C*07:02:80 | 5 | G*01:01:02:01 | 2 |
| C*02:02:01 | H*01:01:01:01 | 8 | C*17:03:01:01 | 5 | H*02:02 | 6 |
| C*02:02:01 | J*01:01:01:01 | 6 | C*17:03:01:01 | 5 | J*02:01 | 2 |
| C*02:02:01 | K*01:01:01:01 | 6 | C*17:03:01:01 | 5 | K*01:02 | 1 |
| C*02:02:01 | L*01:01:01:01 | 5 | C*17:03:01:01 | 5 | L*01:01:02 | 1 |
| C*02:02:01 | Y*01:01 | 8 | C*17:03:01:01 | 5 | Y*02:01 | 0 |

HLA Class I homology for exon 1, Levenshtein distance within and across genes. Exon 1 is usually 73 base pairs long. Distances within genes that are greater than potential distances across genes are highlighted. The full table is symmetric, and for the sake of brevity we exclude the rows where the non-common class I genes (ex. HLA-E) are listed first. There are no interesting cases between these non-common class I genes. Distances calculated from alleles where the full exon sequence is known according to IMGT[17], release 3.30.

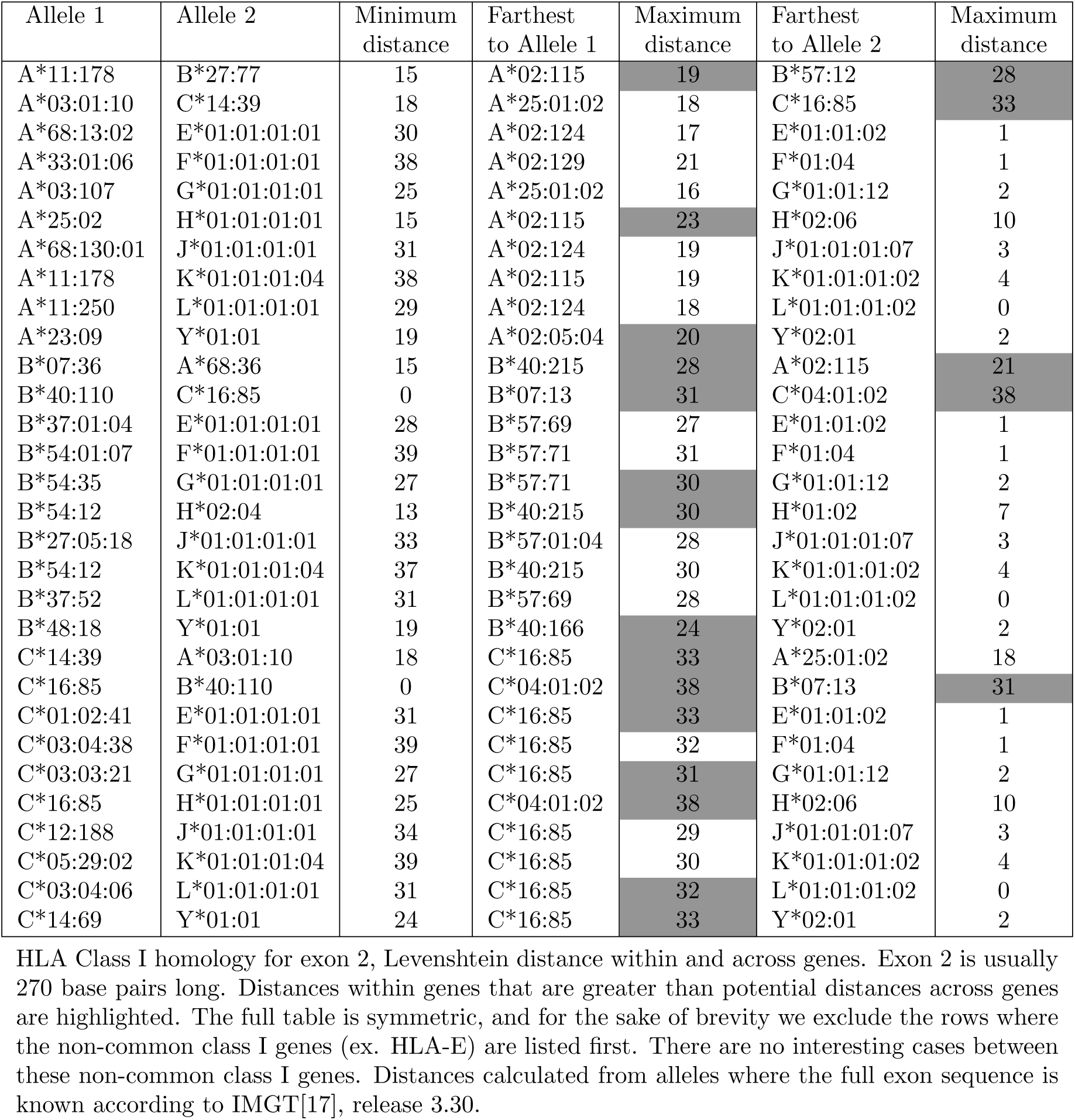

| Allele 1 | Allele 2 | Minimum distance | Farthest to Allele 1 | Maximum distance | Farthest to Allele 2 | Maximum distance |
| --- | --- | --- | --- | --- | --- | --- |
| A*11:178 | B*27:77 | 15 | A*02:115 | 19 | B*57:12 | 28 |
| A*03:01:10 | C*14:39 | 18 | A*25:01:02 | 18 | C*16:85 | 33 |
| A*68:13:02 | E*01:01:01:01 | 30 | A*02:124 | 17 | E*01:01:02 | 1 |
| A*33:01:06 | F*01:01:01:01 | 38 | A*02:129 | 21 | F*01:04 | 1 |
| A*03:107 | G*01:01:01:01 | 25 | A*25:01:02 | 16 | G*01:01:12 | 2 |
| A*25:02 | H*01:01:01:01 | 15 | A*02:115 | 23 | H*02:06 | 10 |
| A*68:130:01 | J*01:01:01:01 | 31 | A*02:124 | 19 | J*01:01:01:07 | 3 |
| A*11:178 | K*01:01:01:04 | 38 | A*02:115 | 19 | K*01:01:01:02 | 4 |
| A*11:250 | L*01:01:01:01 | 29 | A*02:124 | 18 | L*01:01:01:02 | 0 |
| A*23:09 | Y*01:01 | 19 | A*02:05:04 | 20 | Y*02:01 | 2 |
| B*07:36 | A*68:36 | 15 | B*40:215 | 28 | A*02:115 | 21 |
| B*40:110 | C*16:85 | 0 | B*07:13 | 31 | C*04:01:02 | 38 |
| B*37:01:04 | E*01:01:01:01 | 28 | B*57:69 | 27 | E*01:01:02 | 1 |
| B*54:01:07 | F*01:01:01:01 | 39 | B*57:71 | 31 | F*01:04 | 1 |
| B*54:35 | G*01:01:01:01 | 27 | B*57:71 | 30 | G*01:01:12 | 2 |
| B*54:12 | H*02:04 | 13 | B*40:215 | 30 | H*01:02 | 7 |
| B*27:05:18 | J*01:01:01:01 | 33 | B*57:01:04 | 28 | J*01:01:01:07 | 3 |
| B*54:12 | K*01:01:01:04 | 37 | B*40:215 | 30 | K*01:01:01:02 | 4 |
| B*37:52 | L*01:01:01:01 | 31 | B*57:69 | 28 | L*01:01:01:02 | 0 |
| B*48:18 | Y*01:01 | 19 | B*40:166 | 24 | Y*02:01 | 2 |
| C*14:39 | A*03:01:10 | 18 | C*16:85 | 33 | A*25:01:02 | 18 |
| C*16:85 | B*40:110 | 0 | C*04:01:02 | 38 | B*07:13 | 31 |
| C*01:02:41 | E*01:01:01:01 | 31 | C*16:85 | 33 | E*01:01:02 | 1 |
| C*03:04:38 | F*01:01:01:01 | 39 | C*16:85 | 32 | F*01:04 | 1 |
| C*03:03:21 | G*01:01:01:01 | 27 | C*16:85 | 31 | G*01:01:12 | 2 |
| C*16:85 | H*01:01:01:01 | 25 | C*04:01:02 | 38 | H*02:06 | 10 |
| C*12:188 | J*01:01:01:01 | 34 | C*16:85 | 29 | J*01:01:01:07 | 3 |
| C*05:29:02 | K*01:01:01:04 | 39 | C*16:85 | 30 | K*01:01:01:02 | 4 |
| C*03:04:06 | L*01:01:01:01 | 31 | C*16:85 | 32 | L*01:01:01:02 | 0 |
| C*14:69 | Y*01:01 | 24 | C*16:85 | 33 | Y*02:01 | 2 |

HLA Class I homology for exon 2, Levenshtein distance within and across genes. Exon 2 is usually 270 base pairs long. Distances within genes that are greater than potential distances across genes are highlighted. The full table is symmetric, and for the sake of brevity we exclude the rows where the non-common class I genes (ex. HLA-E) are listed first. There are no interesting cases between these non-common class I genes. Distances calculated from alleles where the full exon sequence is known according to IMGT[17], release 3.30.

| Allele 1 | Allele 2 | Minimum distance | Farthest to Allele 1 | Maximum distance | Farthest to Allele 2 | Maximum distance |
| --- | --- | --- | --- | --- | --- | --- |
| A*32:89 | B*27:71 | 9 | A*02:38 | 21 | B*07:78 | 22 |
| A*30:52 | C*12:96 | 13 | A*01:01:31 | 25 | C*17:19 | 17 |
| A*02:594 | E*01:03:01:01 | 44 | A*01:01:31 | 26 | E*01:07 | 2 |
| A*03:186 | F*01:01:01:01 | 44 | A*01:130 | 21 | F*01:01:01:02 | 0 |
| A*24:333 | G*01:20 | 26 | A*26:68 | 24 | G*01:17 | 5 |
| A*30:02:05 | H*02:04 | 12 | A*01:130 | 22 | H*03:01 | 8 |
| A*30:04:01 | J*01:01:01:01 | 31 | A*01:130 | 20 | J*01:01:01:07 | 1 |
| A*30:01:07 | K*01:01:01:01 | 30 | A*01:130 | 24 | K*01:01:01:02 | 3 |
| A*03:186 | L*01:01:01:01 | 38 | A*01:130 | 21 | L*01:02 | 3 |
| A*29:02:16 | Y*01:01 | 7 | A*02:38 | 18 | Y*03:01 | 0 |
| B*27:71 | A*32:89 | 9 | B*07:78 | 22 | A*02:38 | 21 |
| B*35:205 | C*15:117 | 2 | B*08:14 | 20 | C*04:77 | 15 |
| B*14:22 | E*01:03:01:01 | 40 | B*40:298:02 | 23 | E*01:07 | 2 |
| B*27:90:01 | F*01:01:01:01 | 37 | B*07:78 | 21 | F*01:01:01:02 | 0 |
| B*14:33 | G*01:17 | 26 | B*07:78 | 23 | G*01:15 | 5 |
| B*07:225 | H*02:04 | 13 | B*45:14 | 28 | H*03:01 | 8 |
| B*14:33 | J*01:01:01:01 | 24 | B*07:78 | 23 | J*01:01:01:07 | 1 |
| B*15:323 | K*01:01:01:01 | 35 | B*45:03 | 21 | K*01:01:01:02 | 3 |
| B*14:02:10 | L*01:01:01:01 | 34 | B*40:298:02 | 23 | L*01:02 | 3 |
| B*47:10 | Y*01:01 | 15 | B*08:84 | 22 | Y*03:01 | 0 |
| C*12:96 | A*30:52 | 13 | C*17:19 | 17 | A*01:01:31 | 25 |
| C*15:111 | B*40:120 | 2 | C*03:296 | 15 | B*08:14 | 19 |
| C*16:02:05 | E*01:03:01:01 | 36 | C*17:19 | 16 | E*01:07 | 2 |
| C*07:140 | F*01:01:01:01 | 37 | C*03:04:18 | 18 | F*01:01:01:02 | 0 |
| C*01:02:32 | G*01:20 | 21 | C*03:234 | 16 | G*01:17 | 5 |
| C*17:29 | H*02:04 | 17 | C*12:183 | 18 | H*03:01 | 8 |
| C*06:103 | J*01:01:01:01 | 19 | C*17:19 | 18 | J*01:01:01:07 | 1 |
| C*06:153 | K*01:01:01:01 | 36 | C*03:287 | 14 | K*01:01:01:02 | 3 |
| C*07:12 | L*01:01:01:01 | 32 | C*03:04:18 | 17 | L*01:02 | 3 |
| C*02:07 | Y*01:01 | 18 | C*03:234 | 18 | Y*03:01 | 0 |

HLA Class I homology for exon 3, Levenshtein distance within and across genes. Exon 3 is usually 276 base pairs long. Distances within genes that are greater than potential distances across genes are highlighted. The full table is symmetric, and for the sake of brevity we exclude the rows where the non-common class I genes (ex. HLA-E) are listed first. There are no interesting cases between these non-common class I genes. Distances calculated from alleles where the full exon sequence is known according to IMGT[17], release 3.30.

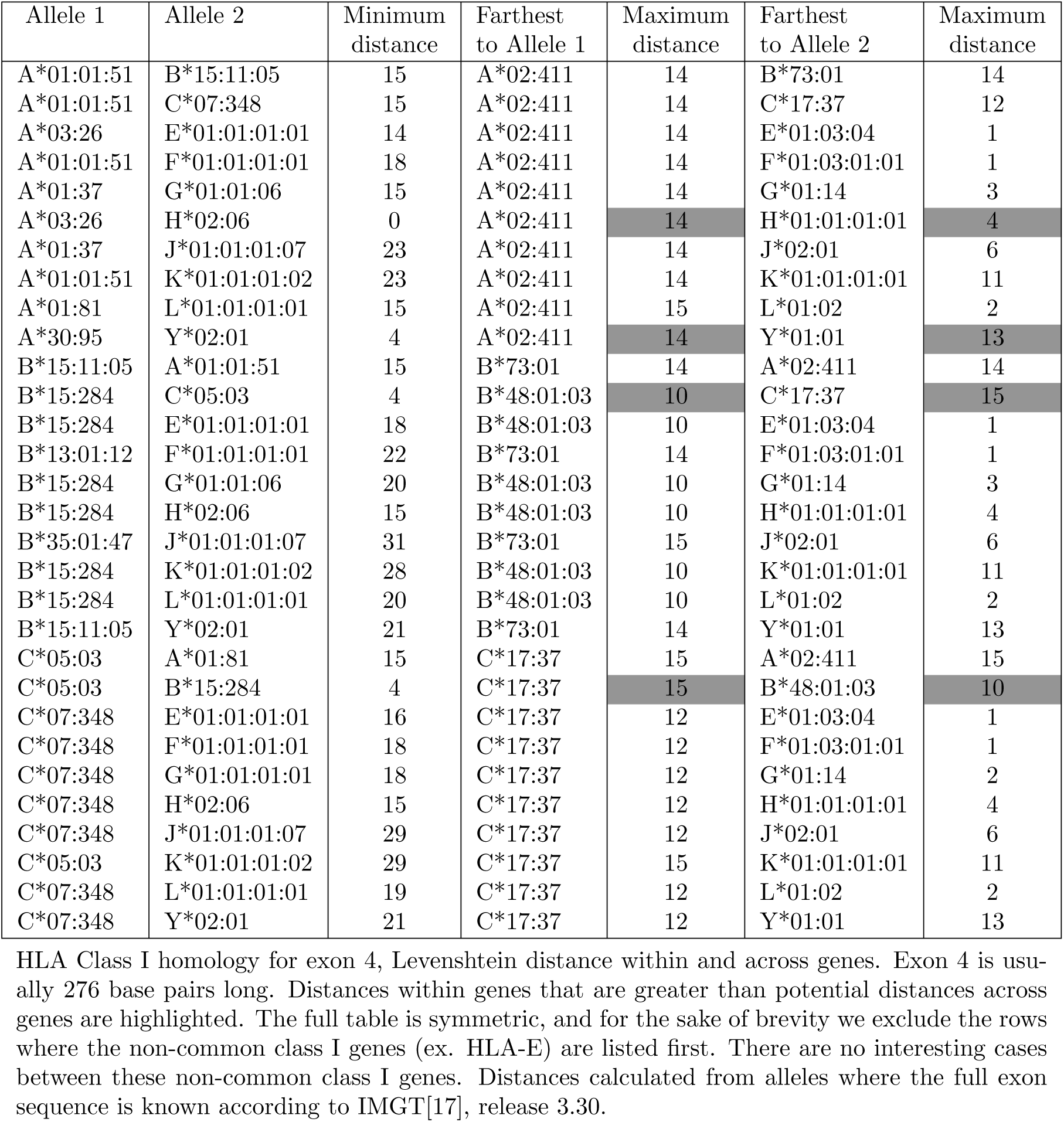

| Allele 1 | Allele 2 | Minimum distance | Farthest to Allele 1 | Maximum distance | Farthest to Allele 2 | Maximum distance |
| --- | --- | --- | --- | --- | --- | --- |
| A*01:01:51 | B*15:11:05 | 15 | A*02:411 | 14 | B*73:01 | 14 |
| A*01:01:51 | C*07:348 | 15 | A*02:411 | 14 | C*17:37 | 12 |
| A*03:26 | E*01:01:01:01 | 14 | A*02:411 | 14 | E*01:03:04 | 1 |
| A*01:01:51 | F*01:01:01:01 | 18 | A*02:411 | 14 | F*01:03:01:01 | 1 |
| A*01:37 | G*01:01:06 | 15 | A*02:411 | 14 | G*01:14 | 3 |
| A*03:26 | H*02:06 | 0 | A*02:411 | 14 | H*01:01:01:01 | 4 |
| A*01:37 | J*01:01:01:07 | 23 | A*02:411 | 14 | J*02:01 | 6 |
| A*01:01:51 | K*01:01:01:02 | 23 | A*02:411 | 14 | K*01:01:01:01 | 11 |
| A*01:81 | L*01:01:01:01 | 15 | A*02:411 | 15 | L*01:02 | 2 |
| A*30:95 | Y*02:01 | 4 | A*02:411 | 14 | Y*01:01 | 13 |
| B*15:11:05 | A*01:01:51 | 15 | B*73:01 | 14 | A*02:411 | 14 |
| B*15:284 | C*05:03 | 4 | B*48:01:03 | 10 | C*17:37 | 15 |
| B*15:284 | E*01:01:01:01 | 18 | B*48:01:03 | 10 | E*01:03:04 | 1 |
| B*13:01:12 | F*01:01:01:01 | 22 | B*73:01 | 14 | F*01:03:01:01 | 1 |
| B*15:284 | G*01:01:06 | 20 | B*48:01:03 | 10 | G*01:14 | 3 |
| B*15:284 | H*02:06 | 15 | B*48:01:03 | 10 | H*01:01:01:01 | 4 |
| B*35:01:47 | J*01:01:01:07 | 31 | B*73:01 | 15 | J*02:01 | 6 |
| B*15:284 | K*01:01:01:02 | 28 | B*48:01:03 | 10 | K*01:01:01:01 | 11 |
| B*15:284 | L*01:01:01:01 | 20 | B*48:01:03 | 10 | L*01:02 | 2 |
| B*15:11:05 | Y*02:01 | 21 | B*73:01 | 14 | Y*01:01 | 13 |
| C*05:03 | A*01:81 | 15 | C*17:37 | 15 | A*02:411 | 15 |
| C*05:03 | B*15:284 | 4 | C*17:37 | 15 | B*48:01:03 | 10 |
| C*07:348 | E*01:01:01:01 | 16 | C*17:37 | 12 | E*01:03:04 | 1 |
| C*07:348 | F*01:01:01:01 | 18 | C*17:37 | 12 | F*01:03:01:01 | 1 |
| C*07:348 | G*01:01:01:01 | 18 | C*17:37 | 12 | G*01:14 | 2 |
| C*07:348 | H*02:06 | 15 | C*17:37 | 12 | H*01:01:01:01 | 4 |
| C*07:348 | J*01:01:01:07 | 29 | C*17:37 | 12 | J*02:01 | 6 |
| C*05:03 | K*01:01:01:02 | 29 | C*17:37 | 15 | K*01:01:01:01 | 11 |
| C*07:348 | L*01:01:01:01 | 19 | C*17:37 | 12 | L*01:02 | 2 |
| C*07:348 | Y*02:01 | 21 | C*17:37 | 12 | Y*01:01 | 13 |

HLA Class I homology for exon 4, Levenshtein distance within and across genes. Exon 4 is usually 276 base pairs long. Distances within genes that are greater than potential distances across genes are highlighted. The full table is symmetric, and for the sake of brevity we exclude the rows where the non-common class I genes (ex. HLA-E) are listed first. There are no interesting cases between these non-common class I genes. Distances calculated from alleles where the full exon sequence is known according to IMGT[17], release 3.30.

## B Partition Maps

In order to achieve an efficient forward pass we need to make the recursive calls

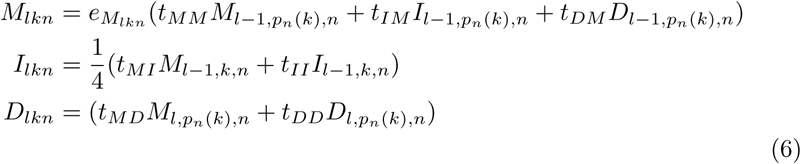

as fast as possible. This requires a way of storing and updating per-allele likelihood values. The general data structure that we seek is map from alleles to values. Naively storing a value for each allele is wasteful and slow as discussed in section 2.3.2, so a linear data structure such as an array or list is not suitable.

We know that we want a mapping where the number of key-value pairs is on the order of the number of values *(v)*, as opposed to the number of keys. Therefore, how do we aggregate these keys, or is there an efficient representations of a large set of alleles into one key?

The size *(N)* of the full set of alleles is specified ahead of time^20^, The desired set of keys always form partition elements of this set. This is the origin of the name partition map. The problem that remains is to determine an efficient representation for these sets.

One can use a bitvector for the keys. An association list of bitvector and unique values. A bitvector represents an allele’s presence in a subset and is *N*/64 words long. This is fairly efficient (as compared to naively using maps or arrays) as computing intersections and set-differences can be accomplished with fast bit operations. This approach results in a running time that is approximately 20% of the naive, “for-each-allele” approach. The main drawback with this approach is that the resulting representation is inefficient for small subsets (e.g. a single allele). Furthermore, it doesn’t actually achieve the reduced theoretical running time that we are after *N*/64 = *O*(*N*); during each of the comparisons over the keys of this operation, we are still imposing this cost and it never shrinks.

An (slight) improvement can be achieved with if we store the allele subsets as interval pairs of their indices: (1, 4) would mean a set from the first to fourth allele. Now we can impose a natural, ascending, order on the association list; compare by the first element of the pair. By maintaining the list in ascending order; this guarantee’s that there is *always* an intersection at the first (head) elements of the two lists during merging, and after computing a new value there is less of the list to process. Unfortunately, the full size of this list is still not effectively constrained as we can have the same value, at later alleles.

The final improvement, the one that is used in production, is to use a list of pairs as the keys, one associated with a unique value. Two ordering are imposed, the key sub-lists are in ascending order, and their first elements are also in ascending order. Unfortunately, when constructing these lists we have to traverse the accumulator after each comparison looking for values that are equal. This turns out to be worth the extra cost as it effectively compresses the final association list.

## Footnotes

1 There is, thankfully, no HLA-“I” gene.

2 HLA-B is missing the last 8th exon which is 5 bp long.

3 The methods described here are easily generalized to different ploidies.

4 Chimeric reads are very rare and we will ignore them for the sake of this analysis.

5 A full Bayesian framework would infer posterior probabilities for these values. We do not do so because it would greatly complicate inference and knowing these values is (probably) of limited importance to HLA typing.

6 A polymerase for example.

7 Go backwards. Move to a different reference position.

8 In reality alleles have different lengths due to inserts and deletions. Let *K* represent some summary statistic such as mean or max of all allele lengths in the analysis.

9 Depends upon where the read starts.

10 Assuming a flat prior.

11 Out of 149 total, this was an exome sequenced sample.

12 For the reads where this locus had the maximum likelihood.

13 Specifically, a triplet of pairs since we don’t consider non-diploid possibilities such as 6 HLA-A’s.

14 As previously mentioned, we will leave aside the problem of detecting new HLA alleles. Our oracle only specifies alleles within our known universe.

15 Greater than 1 ^6^.

16 Not part of this G group.

17 This base had a Q16 score.

18 Not part of this G group, even though at the time of writing IMGT does not acknowledge a A*26:01:01G group. A*26:08 differs by 2 consecutive bases within the third exon with respect to any allele starting with A*26:01:01.

19 Greater than 10^6^ times.

20 When we specify the global alignment file from IMGT, or merge gDNA and cDNA data.

